# The VTA-BLA-NAc circuit for sex reward inhibited by VTA GABAergic neurons under stress in male mice

**DOI:** 10.1101/2021.01.20.427537

**Authors:** Linshan Sun, Jingjing You, Minghu Cui, Fengjiao Sun, Jiangong Wang, Wentao Wang, Dan Wang, Dunjiang Liu, Zhicheng Xu, Changyun Qiu, Bin Liu, Haijing Yan

## Abstract

Anhedonia, inability to experience pleasure from rewarding or enjoyable activities, is the prominent symptom of depression that involves dysfunction of the reward processing system. Both genetic predisposition and life events are thought to increase the risk for depression, in particular life stress. The cellular mechanism underlying stress modulating the reward processing neural circuits and subsequently disrupting reward-related behaviors remains elusive. We identify the VTA-BLA-NAc pathway as being activated by sex reward. Blockade of this circuit induces depressive-like behaviors, while reactivation of VTA neurons associated with sexual rewarding experience acutely ameliorates the impairment of reward-seeking behaviors induced by chronic restraint stress. Our histological and electrophysiological results show that the VTA neuron subpopulation responding to restraint stress inhibits the responsiveness of the VTA dopaminergic neurons to sexual reward. Together, these results reveal the cellular mechanism by which stress influences the brain reward processing system and provide a potential target for depression treatment.

## Introduction

Depression, one of the world’s greatest public health problems, is heterogeneous in aetiology, pathology and treatment responses (Garriock et al, 2010; Fabbri et al, 2017). The existing antidepressant medications, acting on the brain’s serotonergic or noradrenergic systems, usually need at least several weeks to show alleviation effect in depressive symptomatology (Ressler & Nemeroff, 2000; Nestler et al, 2002; Morilak & Frazer, 2004), while approximately 30% of patients show little improvement, who are therefore defined as having treatment-resistant depression (TRD) (Gaynes et al, 2020; Sackeim, 2001; Davidson et al, 2020). Anhedonia, the inability to experience pleasure and insensitivity to naturally rewarding stimuli, is a prominent clinical feature of depression and is considered a potential predictive clinical sign of TRD (McMakin et al, 2012; DeWilde et al, 2015). Accumulating evidence strongly supports dysfunction of the brain reward processing system in depression (Rappaport et al, 2020; Coccurello, 2019).

The mesolimbic system, originating from the ventral tegmental area (VTA) dopaminergic (DA) neurons which project to the nucleus accumbens (NAc), has been extensively involved in regulating motivated behaviors related to reward stimuli and reward-predictive cues (Halbout et al, 2019; Ostlund et al, 2014; Wassum et al, 2013; Yuan et al, 2019), and its abnormalities are associated with Depressive-like behaviors (Krishnan et al, 2007; Chaudhury et al, 2013; Cao et al, 2010). The VTA-NAc pathway does not regulate reward-related behaviors as an independent brain structure, but functions as part of an overlapping and interacting neural circuit. The VTA and NAc receive glutamatergic inputs from the medial prefrontal cortex (mPFC), hippocampus and basolateral amygdala (BLA) (Sesack & Grace, 2010; French & Totterdell, 2003; MacAskill et al, 2012; Britt et al, 2012; Hyman & Malenka, 2001; Nestler & Lüscher, 2019). In return, the VTA neurons also affect the functions of mPFC and hippocampus through axonal innervation (Wittmann et al, 2005; Duszkiewicz et al, 2019; Liu D et al, 2018; Popescu et al, 2016). A surge of research suggests that BLA plays an important role in reward processing, particularly in reward learning and goal-directed behaviors (Kim et al, 2016; Wassum & Izquierdo, 2015). The excitatory transmission from BLA to NAc increases cue-triggered motivated behaviors and supports positive reinforcement (Gore et al, 2015; Di Ciano & Everitt, 2004; Setlow et al, 2002; Stuber et al, 2011). The dopaminergic projections from the VTA to NAc are required for appropriate reward-seeking behaviors regulated by the BLA-NAc pathway (Ambroggi et al, 2008; Stuber et al, 2011). In addition, some neuropharmacological evidence indicates that the VTA may control the activity of BLA-NAc pathway through axonal innervation on BLA neurons (Di Ciano & Everitt, 2004; Lintas et al, 2011). Neverthelss, further research, especially anatomical evidence, is needed to elucidate the neural circuit.

The VTA, a hub of the mesolimbic system that serves an essential role in both reward and aversion (Koob & Le Moal, 2001; de Jong et al, 2019; Russo & Nestler, 2013; Watabe-Uchida et al, 2017), is a heterogeneous brain structure containing dopaminergic (65%), GABAergic (30%) and glutamatergic (5%) neurons (Margolis et al, 2006; Nair-Roberts et al, 2008). The VTA dopaminergic neurons, the primary focus of research on this brain region, have been involved in not only processing rewards and reward-predictive cues (Schultz, 2006; Bayer & Glimcher, 2005), but also responding to aversive and alerting events (de Jong et al, 2019; Bromberg-Martin et al, 2010), and abnormalities in the function of VTA dopaminergic neurons are linked to several neuropsychiatric disorders, including addiction, schizophrenia and depression (Chaudhury et al, 2013; Willuhn et al, 2010; Guillin et al, 2007; Dunlop & Nemeroff, 2007; Nestler & Carlezon, 2006). It is well established that the VTA dopaminergic neurons exhibit rapid and brief burst firing in response to unexpected rewarding stimuli or reward-cues (Watabe-Uchida et al, 2017; Pignatelli & Bonci, 2015). Some studies suggest that the VTA dopaminergic neurons are inhibited by the aversive events (Ungless et al, 2004; Matsumoto & Hikosaka, 2009), while there are paradoxical evidence showing that the VTA dopaminergic neurons are also activated by aversive stimuli (Brischoux et al, 2009; Budygin et al, 2012). Recent research has revealed that the VTA GABAergic neurons are also involved in mediating both reward and aversion (Tan et al, 2012; van Zessen et al, 2012), and are strongly modulated by stress (Tan et al, 2012; Ostroumov et al, 2016), which indicating a potential role in stress-related neuropsychiatric disorders such as depression and post-traumatic stress disorder (PTSD). Research data have shown that VTA GABAergic neurons synapse onto local VTA dopaminergic neurons and exhibit inhibitory effects on addiction and aversion (Matsui et al, 2014; Matsui et al, 2011; Polter et al, 2018; Tan et al, 2012). Despite these knowledge, the role of VTA neurons in normal reward-related behaviors, such as food and sex, remains to be determined, which may help us to further understand the mechanism by which stress induces anhedonia.

## Results

### Brain areas activated during positive experience

To identify brain regions that are activated during a sexual rewarding experience, we caged male mice with female mice for 2 hours (hereafter referred to as a ‘positive experience’) and then performed brain-wide immunofluorescent staining of c-Fos. The histological data showed that there was c-Fos expression in several brain areas, including the basolateral amygdala (BLA), nucleus accumbens (NAc), medial prefrontal cortex (mPFC), ventral tegmental area (VTA), substantia nigra (SN), interpeduncular nucleus (IPN), and dentate gyrus (DG), which all have been implicated in reward processing and motivated behaviors (Zhang et al, 2020; LeGates et al, 2018; Ferenczi et al, 2016; Lammel et al, 2012; Ilango et al, 2014; McLaughlin et al, 2017; Ramirez et al, 2015), in both neutral experience and positive experience mice (Fig 1A-1E’). Positive experience mice exhibited more c-Fos-expressing cells in the BLA, NAc and VTA (Fig 1F, 1G and 1I), but not in the mPFC, SN, IPN or DG (Fig 1H, 1J, 1K and 1L). These results suggest that VTA, BLA and NAc are activated by positive experience and may be involved in processing the sexual reward.

**Figure 1.**
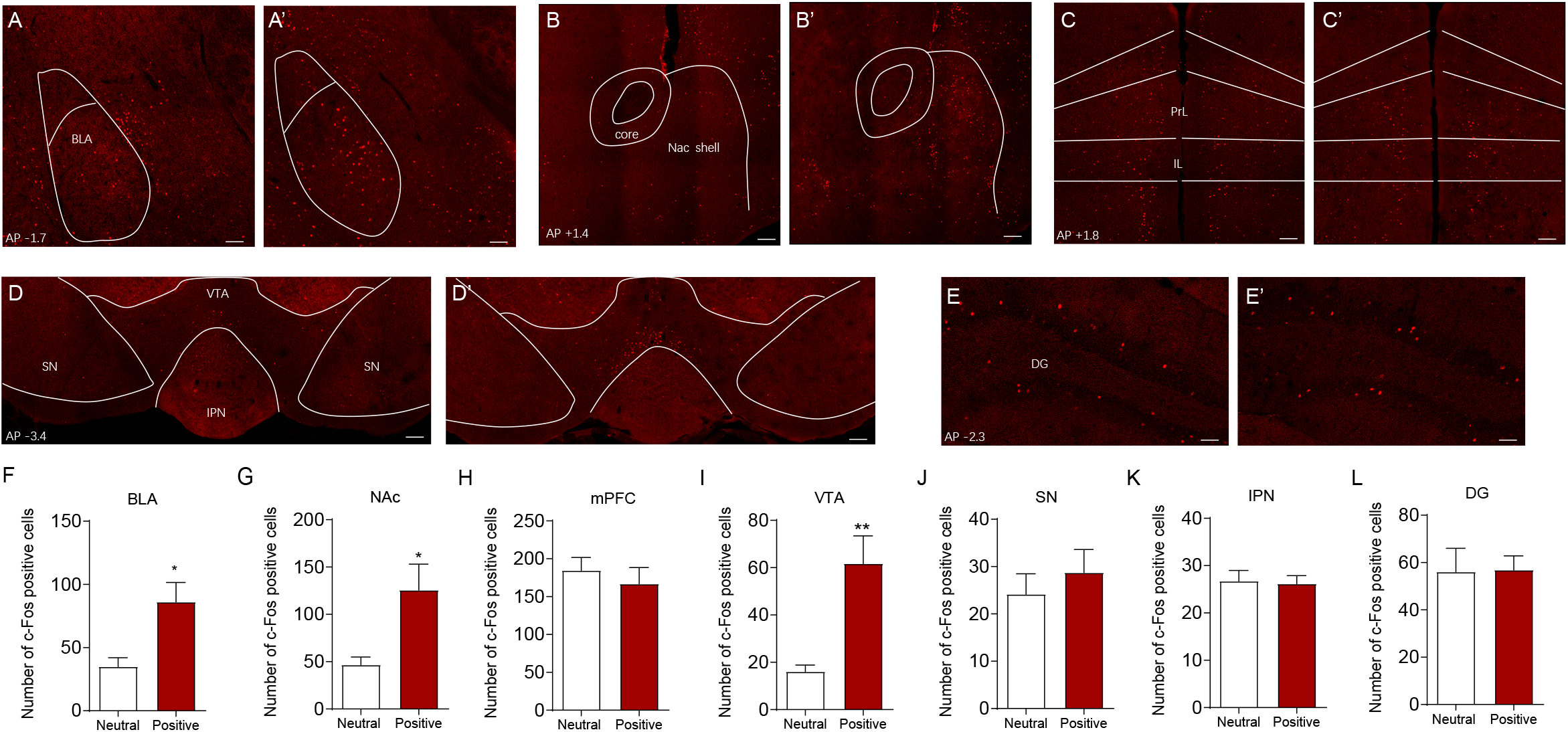
Positive experience increases c-Fos expression in basolateral amygdala, nucleus accumbens and ventral tegmental area. Whole brain c-Fos staining shows that positive experience (A’-E’), but not neutral experience (A-E), elicits increase of c-Fos expression in basolateral amygdala (BLA) (A, A’), nucleus accumbens (NAc) (B, B’), ventral tegmental area (VTA) (D, D’), but not in the medial prefrontal cortex (mPFC, including prelimbic (PrL) and infralimbic (IL)) (C, C’), substantia nigra (SN) (D, D’), interpeduncular nucleus (IPN) (D, D’) or dentate gyrus (DG) (E, E’). Statistical analysis of the histological data revealed a significant increase of c-Fos expression in basolateral amygdala (F, two-tailed unpaired t test, t_(12)_ = 2.993, P = 0.0112), nucleus accumbens (G, two-tailed unpaired t test with Welch’s correction, t_(7.109)_ = 2.747, P = 0.0282) and ventral tegmental area (I, two-tailed unpaired t test with Welch’s correction, t_(6.687)_ = 3.788, P = 0.0074). Neutral experience group n=7; positive experience group n=7. *P<0.05, **P<0.01. Data are presented as means +/− SEM. Annotation (AP): distance from the bregma (mm). Scale bars correspond to 100μm.

### Reactivation of hM3D-labeled VTA neurons during positive experience increases c-Fos expression in BLA and NAc

The VTA plays a central role in reward processing and motivated behaviors through diverse projections to target brain regions, including BLA, NAc and mPFC (Lintas et al, 2011; Beier et al, 2015; Pignatelli & Bonci, 2018; Heymann et al, 2020; Hauser et al, 2017; Kumar et al, 2018; Pessiglione et al, 2006). The BLA has a crucial role in cue-triggered motivated behaviors and its glutamatergic inputs to the NAc has been implicated in reward-seeking behaviors (Ambroggi et al, 2008; Di Ciano & Everitt, 2004; Stuber et al, 2011). A previous pharmacology study indicated that the VTA-BLA-NAc circuit was involved in opiate-related reward processing, but did not provide direct clear anatomical evidence for this circuit (Lintas et al, 2011). We therefore investigated whether the VTA–BLA–NAc circuit was indeed activated during the positive experience. To address this issue, we injected AAV-DIO-hM3D-mCherry into the VTA of Fos-CreER^T2^ mice. After exposure to conspecific females, Fos-CreER^T2^ male mice were intraperitoneally injected with 4-hydroxytamoxifen (4-OHT, 50 mg/kg) to induce the expression of hM3D-mCherry to label VTA-activated neurons during that positive experience. Three weeks later, we intraperitoneally injected clozapine-N-oxide (CNO,0.3mg/kg) to activate previous hM3D-mCherry-labeled VTA neurons (Fig 2A). Consistent with increase of c-Fos expression in VTA (Fig 1D, 1D’ and 1I), positive experience induced more hM3D-mCherry-labeled VTA neurons (Fig 2B, 2B’ and 2I). Consequently, there were more c-Fos-positive VTA neurons after intraperitoneal injection of CNO (Fig 2C, 2C’ and 2I’). Previous studies showed that VTA neurons could be excited by both rewarding and aversive stimuli (Lintas et al, 2011; Beier et al, 2015; Pignatelli & Bonci, 2018; Heymann et al, 2020). As there might be some alerting cues and aversive stimuli throughout the operation of CNO injection, such as catching mice and intraperitoneal injection, we gently handled mice to reduce aversive stimuli. Our data showed that the vast majority of the c-Fos-positive VTA neurons were hM3D-mCherry-labeled in both groups (Fig 2D, 2D’ and 2I’’, neutral 81.53±5.510%, positive 86.41±1.888%), suggesting that the VTA c-Fos expression was induced predominantly through activating hM3D by CNO. We next checked the c-Fos expression in other brain areas that were activated during the positive experience (Fig 1). Reactivation of VTA neurons labelled by positive experience correlated with increase of c-Fos expression in BLA (Fig 2E, 2E’ and 2L) and NAc (Fig 2F, 2F’ and 2M), but not in SN (Figu 2C, 2C’ and 2J), IPN (Fig 2C, 2C’ and 2K), mPFC (Fig 2G, 2G’ and 2N), or DG (Fig 2H, 2H’ and 2O). Together, these results suggest that VTA may be upstream of BLA and NAc during the positive experience.

**Figure 2.**
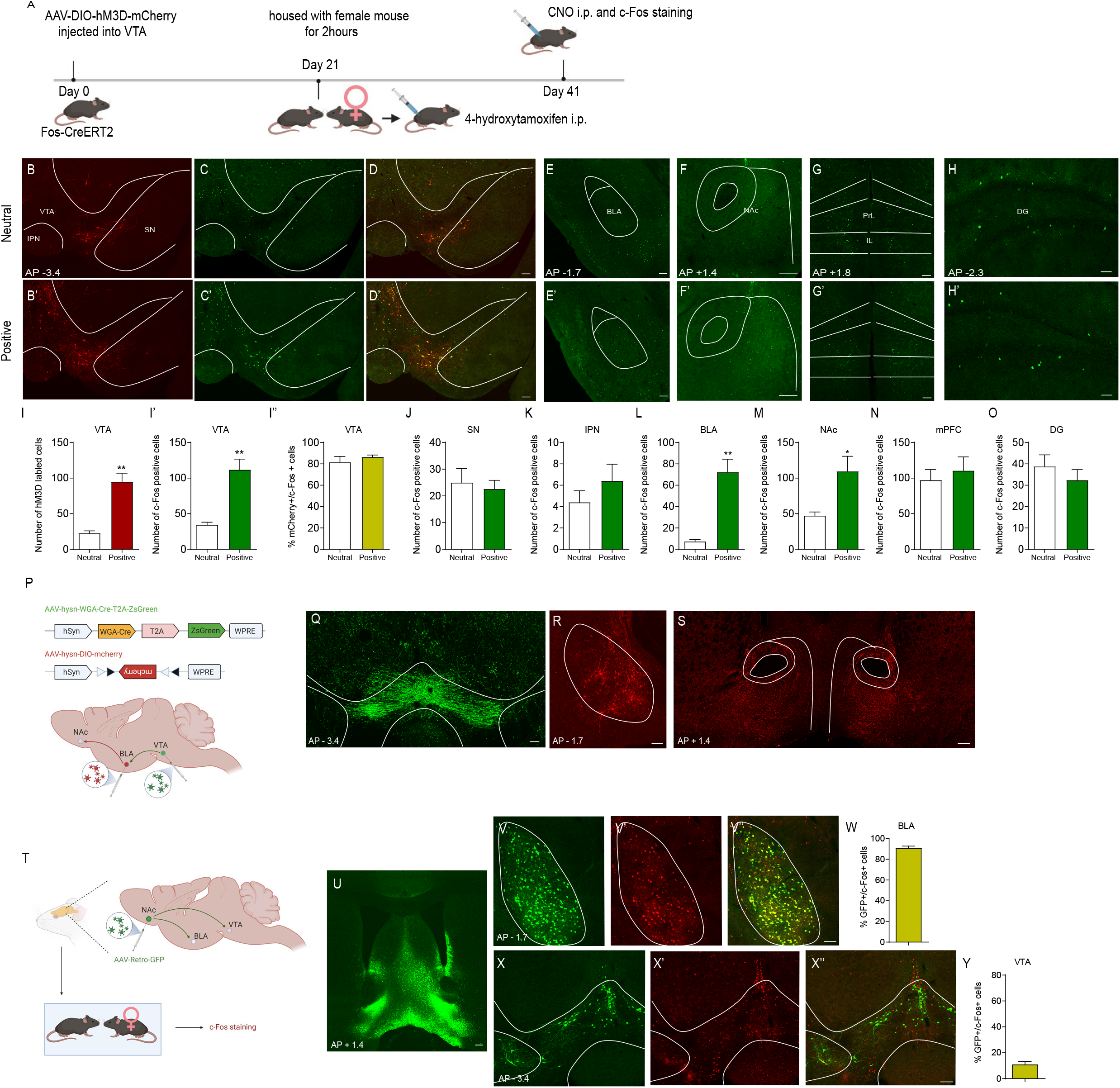
Positive experience activates the VTA-BLA-NAc circuit. (A) The experimental timeline. Intraperitoneal injection of 4-OHT induced hM3D-mCherry labeling in activated cells during neutral experience (B) or positive experience (B’) in the VTA, SN and IPN of Fos-CreERT2 mice. Intraperitoneal injection of clozapine-N-oxide (CNO) activated c-Fos expression in hM3D-mCherry-labeled neurons (neutral experience group, C and D; positive experience group, C’ and D’). C-Fos expression in BLA (E, E’), NAc (F, F’), mPFC (G, G’) and DG (H, H’) after reactivating hM3D-mCherry-labeled VTA neurons. Statistical analysis of the histological data revealed that positive experience induced more hM3D-mCherry-labeled VTA neurons compared with neutral experience (I, two-tailed unpaired t test with Welch’s correction, t_(4.607)_ = 5.852, P = 0.0027), and that the c-Fos positive VTA cells of positive experience group after CNO injection were more than those of neutral experience group (I’, Mann-Whitney U test, P=0.0079) although most VTA cells were hM3D-mCherry-labeled in both groups (I’’, neutral 81.53±5.510%, positive 86.41±1.888%). Few cells were labelled with hM3D-mCherry (B, B’) and only a small number of cells were c-Fos-positive after CNO injection (C, C’) in SN and IPN in both groups. Statistical analysis revealed no significant difference in the number of c-Fos-positive cells in SN (J) and IPN (K) between neutral and positive experience groups. The number of c-Fos-positive cells after clozapine-N-oxide injection was significantly different in BLA (L, two-tailed unpaired t test with Welch’s correction, t_(4.159)_ = 5.265, P = 0.0056) and NAc (M, two-tailed unpaired t test with Welch’s correction, t_(4.461)_ = 2.870, P = 0.04) between neutral and positive experience groups, but not in mPFC (N, Mann-Whitney U test, P=0.5476) and DG (O, two-tailed unpaired t test, t_(8)_ = 0.8750, P = 0.4071). Neutral experience group: n=5; positive experience group: n=5. (P) Diagram illustrating virus injection in target areas. (Q) Representative coronal slice showing the expression of AAV-hSyn-WGA-Cre-T2A-ZsGreen (green) 3 weeks after virus injection into the VTA. (R) Representative coronal slice showing the expression of AAV-CAG-DIO-mCherry (red) 3 weeks after virus injection into the BLA. (S) Representative coronal slice showing the strong mCherry–positive (red) fibers in NAc 3 weeks after virus injection into the BLA. (T) Diagram illustrating virus injection in target areas and subsequent positive experience. (U) Representative image showing the injection sites in the NAc. Representative images showing the expression of GFP (V) and c-Fos (V’) in BLA. (V’’) Representative image showing c-Fos expression in BLA merged with GFP. (W) Statistical analysis showing the percentage of c-Fos positive BLA neurons that were also GFP labeled (69.81±1.731%, n=4). Representative images showing the expression of GFP (X) and c-Fos staining (X’) in NAc. (X’’) Representative merged images. (N) Statistical analysis showing the percentage of c-Fos positive VTA neurons that were also GFP labeled (14.25±3.080%, n=4). *P<0.05, **P<0.01. Data are presented as means +/− SEM. Annotation (AP): distance from the bregma (mm). Scale bars correspond to 100μm.

### VTA-BLA-NAc pathway is activated by positive experience

To verify the hypothetical circuit architecture, i.e. neurons in VTA synapse onto BLA neurons projecting to specific neurons in NAc, we injected AAV-WGA-Cre-T2A-ZsGreen and AAV-DIO-mCherry into VTA and BLA respectively (Fig 2P). The AAV-WGA-Cre-T2A-ZsGreen virus contains transsynaptic tracer wheat-germ agglutinin (WGA) fused to Cre-recombinase, a ZsGreen reporter, and a linker peptide T2A (2A peptide derived from insect Thosea asigna virus) whose “self-cleaving” would generate two proteins, WGA-Cre and ZsGreen (Hadpech et al, 2018). ZsGreen would label the virus infected VTA neurons (Fig 2Q) and WGA-Cre would be released into synaptic cleft and taken up by the adjacent neuron (Libbrecht et al, 2017). As WGA-Cre could undergo both anterograde and retrograde transneuronal transfer (Yoshihara et al, 1999; Horowitz et al, 1999), WGA-Cre could enter BLA neurons projecting to or receiving inputs from VTA, and induce mCherry expression in the AAV-DIO-mCherry-infected BLA neurons (Fig 2R). We observed strong mCherry positive fibers in NAc (Fig 2S) that should be derived from mCherry-labled BLA neurons receiving inputs from VTA. These results supported the hypothetical architecture of neural circuit that neurons in BLA synapsing onto NAc neurons received inputs from VTA neurons, and indicated that reactivation of VTA neurons labelled by positive experience may increase c-Fos expression in BLA and VTA through the VTA-BLA-NAc pathway, but did not rule out the possibility that increasing c-Fos expression through VTA-BLA or VTA-NAc pathways.

To address this issue, we performed the pseudotyped rabies virus (RABV)-based monosynaptic retrograde tracing (Tervo et al, 2016; Wickersham et al, 2007). The AAV-Retro-GFP was injected into NAc (Fig 2U) to label projection neurons from BLA and VTA (Fig 2V and 2X). C-Fos staining revealed activated neurons in BLA and NAc during positive experience (Fig 2V’ and 2X’). The c-Fos positive BLA neurons were mostly GFP-labelled (Fig 2V’’ and 2W, 90.97±1.862%, n=4), whereas few c-Fos positive VTA neurons were GFP-labelled (Fig 2X’’ and 2Y, 11.03±2.189%, n=4). Collectively, our data showed that most BLA neurons projecting to NAc were activated by positive experience, whereas only a minority of VTA neurons projecting to NAc were activated, suggesting that reactivation of VTA neurons previously activated by positive experience may activate NAc through the VTA-BLA-NAc pathway.

### Blocking the VTA-BLA-NAc circuit induces depressive-like behaviors

It has become clear that dysfunctions of specific brain networks mediating mood and reward signals underly a variety of mood disorders, including depression and anxiety (Nestler, 2015; Lammel et al, 2014; Kaufling et al, 2017; Stamatakis et al, 2014; Lebow & Chen, 2016). To determine whether blocking the VTA-BLA-NAc circuit responding to positive experience would induce depressive-like behaviors, we injected AAV-DIO-hM4D-mCherry (AAV-DIO-mCherry as control) into VTA or BLA and simultaneously implanted bilateral cannula into BLA or NAc of Fos-CreER^T2^ mice. After three weeks, the mice were exposed to conspecific females for two hours and then immediately received intraperitoneal injection of 4-OHT to induce activity-dependent hM4D-mCherry labelling of VTA or BLA neurons activated by the positive experience. Three weeks later, the mice were subjected to behavioral tests while the synaptic communication from VTA to BLA or from BLA to NAc was silenced through bilateral infusion of CNO (900 pmol/0.3μl /side) into BLA or NAc (Stachniak et al, 2014; Rinker et al, 2017; Mahler et al, 2014) (Fig 3A and 3E).

**Figure 3.**
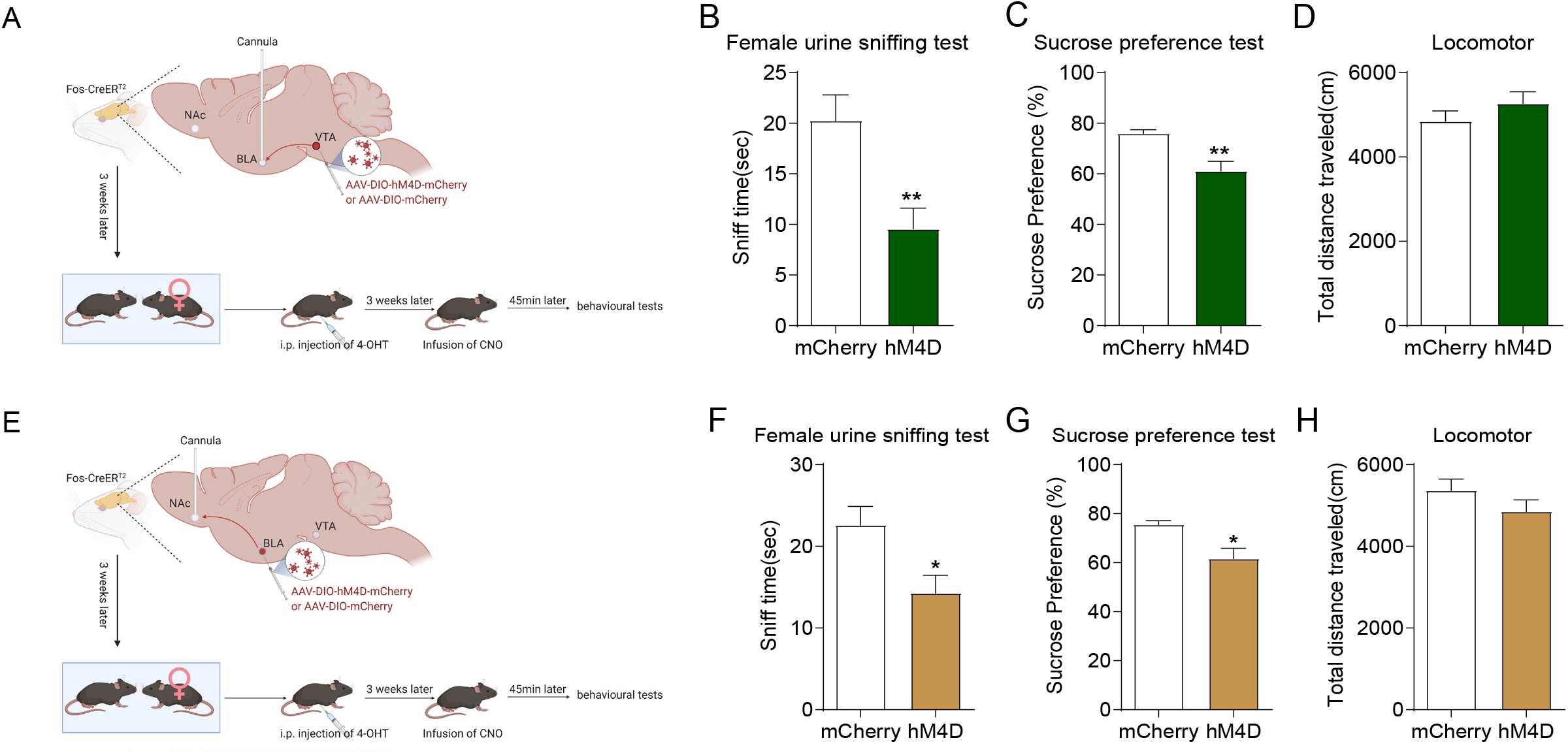
Blockade of VTA-BLA-NAc circuit responding to positive experience induces depressive-like behaviors. (A) Diagram illustrating virus injection in target areas and subsequent experiments. Blocking the projections from VTA to BLA induced depressive-like behaviors, as evaluated by female urine sniffing test (B, Mann-Whitney U test, P=0.0030) and sucrose preference test (C, two-tailed unpaired t test with Welch’s correction, t_(11.73)_=3.584, P = 0.0039), but did not influence the locomotor activity (D, two-tailed unpaired t test, t_(16)_ = 1.088, P = 0.2927). mCherry group: n=8, hM4D group: n=10 in FUST(B), SPT(C) and Locomotor (D). (E) Diagram illustrating virus injection in target areas and subsequent experiments. Blocking the projections from BLA to NAc decreased the sniff time (F, two-tailed unpaired t test, t_(17)_ = 2.627, P = 0.0176) and sucrose preference (G, two-tailed unpaired t test with Welch’s correction, t_(11.41)_=3.024, P = 0.0111), but did not influence the locomotor activity (H, two-tailed unpaired t test, t_(17)_ = 1.088, P = 0.2164). mCherry: group n=9, hM4D group: n=10 in FUST(F), SPT(G) and Locomotor (H). *P<0.05, **P<0.01. Data are presented as means +/− SEM.

We preformed female urine sniff test (FUST) and sucrose preference test (SPT) to evaluate the depression level. Anhedonia is a prominent symptom of depression, which is inability to experience pleasure from previously pleasurable activities, such as sex and food (Coccurello, 2019; Rizvi et al, 2016). FUST and SPT measure anhedonia in mice based on reward-seeking behaviors on female pheromonal odors and sucrose respectively (Malkesman et al, 2010; Liu MY et al, 2018). Our data showed that blocking the VTA-BLA pathway responding to positive experience caused decrease in both sniff time (Fig 3B) and sucrose preference (Fig 3C), but did not affect the locomotor activity (Fig 3D). Similar results were obtained when blocking the BLA-NAc pathway (Fig 3F-H). Collectively, these results showed that blocking the VTA-BLA-NAc circuit responding to positive experience induced depressive-like behaviors, indicating a possibility that reactivating the VTA neurons previously activated by positive experience may ameliorate the depressive symptoms.

### Reactivation of the VTA neurons previously activated by positive experience reverses chronic restraint stress-induced depressive-like behaviors

Previous studies suggest that malfunctions of the brain’s reward circuits play an important role in mediating stress-elicited depression-like behaviors (Lammel et al, 2014; Kaufling et al, 2017). We therefore examined whether reactivation of the VTA neurons previously activated during positive experience could ameliorate depressive-like behaviors induced by chronic restraint stress (Fig 4A). First, Fos-CreER^T2^ mice were injected with AAV-DIO-hM3D-mCherry into the VTA and individually housed until the end of all experiments. Mice were assigned into two groups with no statistical difference in SPT or FUST (Fig 4B and 4C, Basal). Positive experience mice were housed with oestrous female mice for 2 hours, and the neutral experience mice was housed with fake toy mice. Then immediately, all mice received intraperitoneal injection of 4-OHT to allow activity-dependent hM3D-mCherry labelling of VTA neurons activated by the positive experience. The following behavioral experiments were all carried out 45 min after CNO injection. Two weeks later, the SPT showed no difference between the two groups (Fig 4B, Basal-CNO), while the FUST showed significant increase in sniffing time in positive experience mice (Fig 4C, Basal-CNO). After 15 days of restraint stress treatment, positive experience mice exhibited more sucrose preference (Fig 4B, Restraint-CNO) and sniffing time (Fig 4C, Restraint-CNO). Forced swim test (FST) is widely used to assess learned helplessness, another feature of depressive-like behavior, by measuring the immobility time (Cryan & Holmes, 2005). Positive experience mice exhibited reduced immobility time (Fig 4D, left), but no change in the latency to immobility from the start of the test (Fig 4D, right). The total distance travelled suggested that there was no significant difference in locomotor activity between the two groups (Fig 4E). Together, these results suggest that reactivation of VTA neurons responding to positive experience can ameliorate depressive-like behaviors induced by chronic restraint stress.

**Figure 4.**
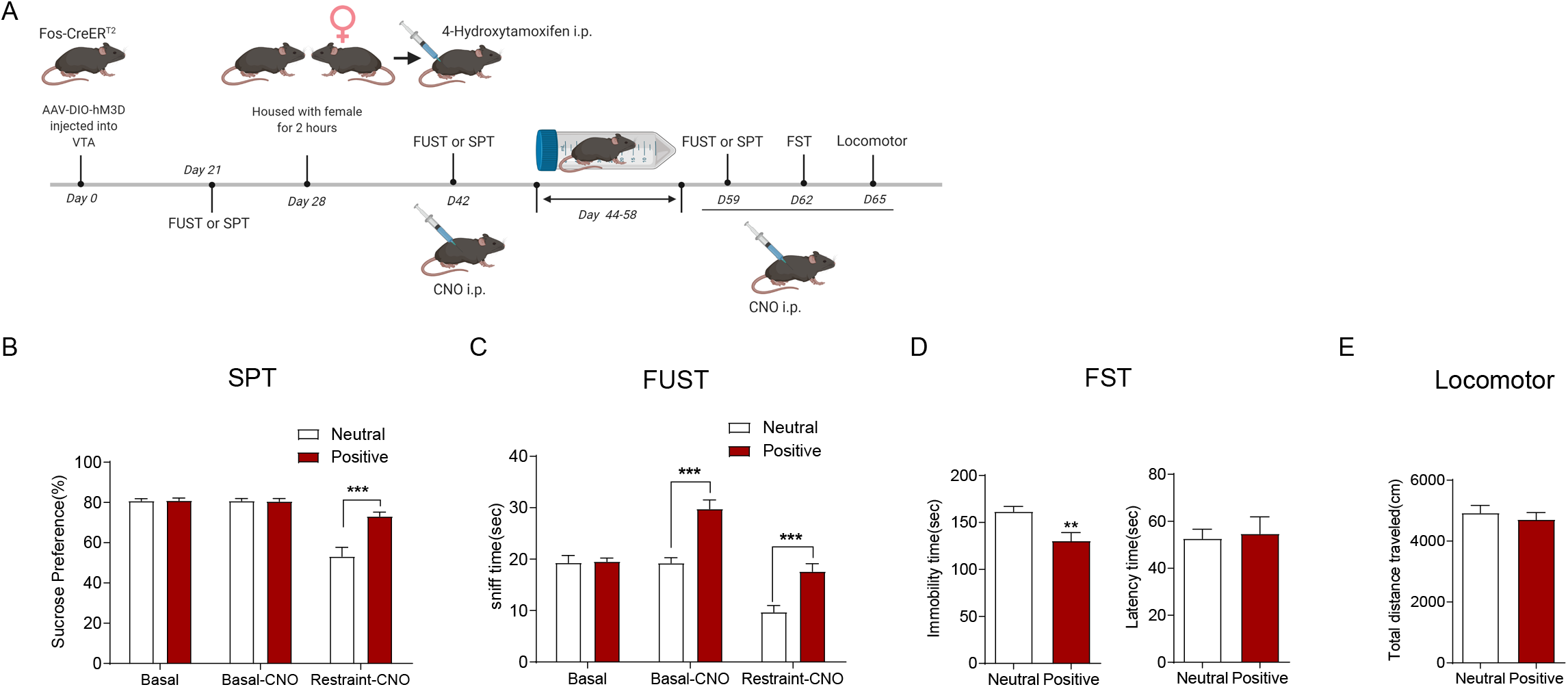
Reactivation of VTA neurons labelled by hm3D during previous positive experience can ameliorate depressive-like behaviors induced by chronic restraint stress. (A) The experimental timeline. (B) Sucrose preference test showing reactivation of hm3D-labeled VTA neurons during previous positive experience by CNO increased sucrose preference under chronic stress condition (subgroups, F(1, 19)=9.634, P=0.0058; treatment, F(2, 38)=53.53, P < 0.001; subgroups X treatment, F(2, 38)=17.30, P<0.001; Neutral VS. Positive, Basal P>0.999, Basal-CNO P=0.9997, Restraint-CNO P<0.0001). (C)FUST showing reactivation of hm3D-labeled VTA neurons during previous positive experience by CNO increased sniff time under chronic stress condition (subgroups, F(1, 21)=37.54, P < 0.001; treatment, F(2,42)=32.22, P<0.001; subgroups X treatment, F(2, 42)=7.818, P=0.0013; Neutral VS. Positive, Basal P=0.9989, Basal-CNO P<0.0001, Restraint-CNO P=0.0002). (D) Forced swim test (FST) showing reactivation of hm3D-labeled VTA neurons during previous positive experience by CNO decreased the immobility time (left, two-tailed unpaired t test, t_(21)_ = 2.911, P = 0.0083), but did not influence the latency time (right, Mann-Whitney U test, P=0.9878), under chronic stress condition. (E) Sucrose preference test showing reactivation of hm3D-labeled VTA neurons during previous positive experience by CNO increased sucrose preference (two-tailed unpaired t test, t_(21)_ = 0.6287, P = 0.5364) under chronic stress condition. Neutral experience group n=10, positive experience group n=11 in SPT (B); Neutral experience group n=11, positive experience group n=12 in FUST (C), FST (D) and locomotor (E). **P<0.01, ***P<0.001. Data are presented as means +/− SEM.

### Distinct subpopulations of VTA neurons activated by positive experience and restraint stress

The VTA is a heterogeneous nucleus including dopaminergic (DAergic), Gamma-aminobutyric acid-ergic (GABAergic), and glutamatergic (Glutergic) neurons, in which DAergic neurons predominate, making up about 55%-65% of the total neurons (Margolis et al, 2006; Tan et al, 2012; Morales & Margolis, 2017). VTA dopaminergic neurons respond not only to rewards and reward-predicting stimuli, but also to aversion, alerting events and behavioral choices (Bayer & Glimcher, 2005; Brischoux et al, 2009; Dautan et al, 2016; Zhou et al, 2019; Howard et al, 2017). This functional heterogeneity is reflected in the anatomically heterogeneous dopaminergic subpopulations connecting with different brain regions (Bromberg-Martin et al, 2010; Engelhard et al, 2019). To clarify the underlying mechanism by which reactivation of VTA neurons activated by positive experience ameliorates the chronic restraint stress-induced depression, we first identified the neuron subpopulation activated during positive experience. We performed c-Fos staining after positive experience in DAT-IRES-Cre;Ai14 mice (Fig 5I), in which tdTomato was exclusively colocalized with the dopaminergic neuron marker TH (97.67±0.7862%, n=4) (Fig 5A-5H), confirming DAT-IRES-Cre-mediated recombination was dopaminergic neuron-specific. Our results showed that the VTA c-Fos positive neurons labelled by positive experience mostly colocalized with tdTomato in the DAT-IRES-Cre;Ai14 mice (68.92±4.902%, n=5) (Fig 5J-5M), suggesting that the neuron subpopulation responding to positive experience is predominantly dopaminergic. Dopaminergic neurons maintain the baseline level of dopamine in downstream neural structures through the tonic firing mode and transit to the phasic burst firing mode to induce a sharp increase in dopamine release in response to both rewarding and aversive/stressful stimuli (Di Ciano & Everitt, 2004; Stuber et al, 2011). We therefore next performed in vivo extracellular recording to examine the activity of VTA dopaminergic neurons after positive experience (Fig 5N) based on the electrophysiological criteria described in previous reports (Tan et al, 2012; Grace & Bunney, 1983; Ungless & Grace, 2012). Consistent with the c-Fos staining results, positive experience increased the number of spontaneously active dopaminergic neurons in the VTA (Fig 5O). The firing rate of spontaneously active dopaminergic neurons was unaffected (Fig 5P), but the percentage of burst firing significantly increased (Fig 5Q). Collectively, these results suggest that the subpopulation of VTA neurons activated during positive experience are dopaminergic.

**Figure 5.**
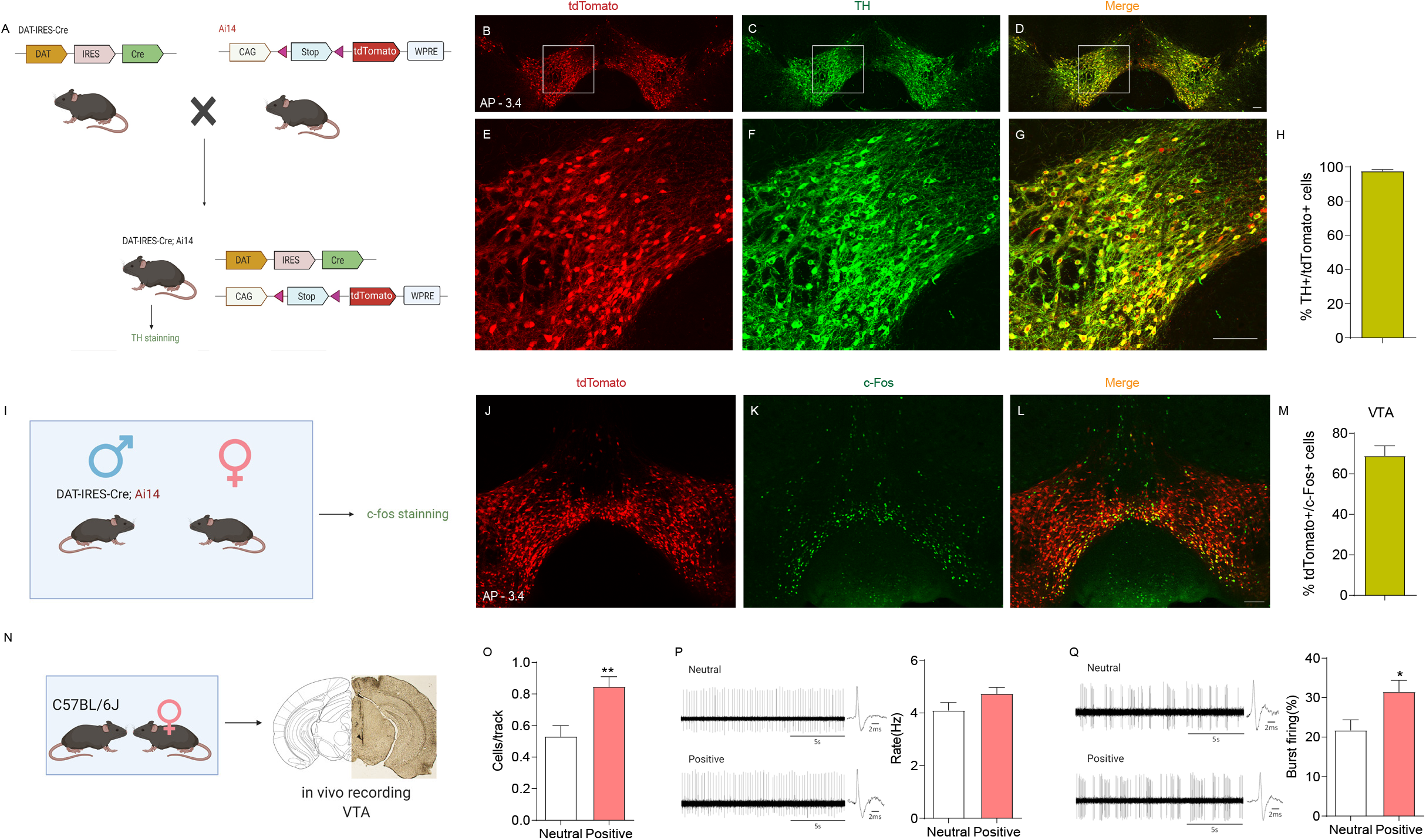
The activated VTA neurons during positive experience are mostly dopaminergic. (A) Schematic of the breeding strategy used to generate mice expressing tdTomato specifically in dopaminergic neurons (DAT-IRES-Cre; Ai14). Representative images showing the expression of tdTomato (B, E) and TH staining (C, F) in VTA. (D, G) Images are merged. (H) Statistical analysis showing the percentage of tdTomato-labeled VTA neurons positive also for TH (97.64±0.7862%, n=4). (I) Schematic showing the experimental procedure for DAT-IRES-Cre;Ai14 mice subject to positive experience and subsequent c-Fos staining. (J) Representative image showing the expression of tdTomato in VTA. (K) Representative image showing the expression of c-Fos in VTA. (L) Images of J and K are merged. (M) Statistical analysis showing the percentage of tdTomato-labeled c-Fos-positive VTA neurons (68.92±4.902%, n=5). (N) Diagram showing the experimental procedure and representative image demonstrating the electrode track through the VTA. (O) Positive experience increased the number of spontaneously active dopaminergic neurons per track in the VTA (Mann-Whitney U test, P=0.0061; Neutral n=9 mice, Positive n=8 mice). (P) Left, representative extracellular voltage traces from VTA dopaminergic neurons; Right, statistical result showing the average firing rate of the spontaneously active dopaminergic neurons (two-tailed unpaired t test, t_(100)_ = 1.710, P=0.0904; Neutral n=9 mice (42 neurons), Positive n=8 mice (60 neurons)). (Q) Left, representative extracellular voltage traces showing the burst firing of VTA dopaminergic neurons; Right, statistical result showing the average percentage of burst firing of the active dopaminergic neurons (Mann-Whitney U test, P=0.0447; Neutral n=9 mice (42 neurons); Positive n=8 mice (60 neurons)). *P<0.05, **P<0.01. Data are presented as means +/− SEM. Annotation (AP): distance from the bregma (mm). Scale bars correspond to 100μm.

We next identified the VTA neuron subpopulation responding to restraint stress. As shown in Fig 6A-6J, restraint stress increased c-Fos expression in VTA, BLA and NAc. We then performed c-Fos staining in VTA of DAT-IRES-Cre;Ai14 mice (Fig 6K) and the result showed that few c-Fos-positive neurons were tdTomato-labelled (17.04±3.513%, n=4) (Fig 6L-6O), suggesting that the activated VTA neurons are mostly non-dopaminergic, indicating that the subpopulation of VTA neurons responding to restraint stress is distinct from that of positive experience. To confirm this observation, we detected both restraint stress- and positive experience-activated neurons in similar brain slices of Fos-CreER^T2^ (Fig 6P) and Fos-CreER^T2^; Ai14 mice (Fig 6U). We injected AAV-DIO-GFP into VTA of Fos-CreER^T2^ mice which were subsequently subjected to 2 hours of restraint stress and received 4-OHT injection to allow activity-dependent GFP labelling of VTA neurons responding to restraint stress. Three weeks later, we housed these mice with conspecific females and performed c-Fos staining. We observed that the VTA neurons activated by restraint stress (green, Fig 6Q) and positive experience (red, Fig 6R) were anatomically distinct subpopulations (Fig 6S). Statistical analysis showed that there were few c-Fos-immunopositive VTA neurons labelled by GFP (3.765±1.285%, n=5) (Fig 6T). Similar results were observed in Fos-CreER^T2^; Ai14 mice. The VTA neurons responding to restraint stress (tdTomato, red, Fig 6V) were distinct from those of positive experience (c-Fos, green, Fig 6W) and few c-Fos-immunopositive VTA neurons were labelled by tdTomato (3.098±1.412%, n=5) (Fig 6X). These results together suggest that the subpopulation of VTA neurons activated by restraint stress is distinct from that activated by positive experience, raising the question of how the VTA neuron subpopulation responding to restraint stress affects the reward-related behaviors.

**Figure 6.**
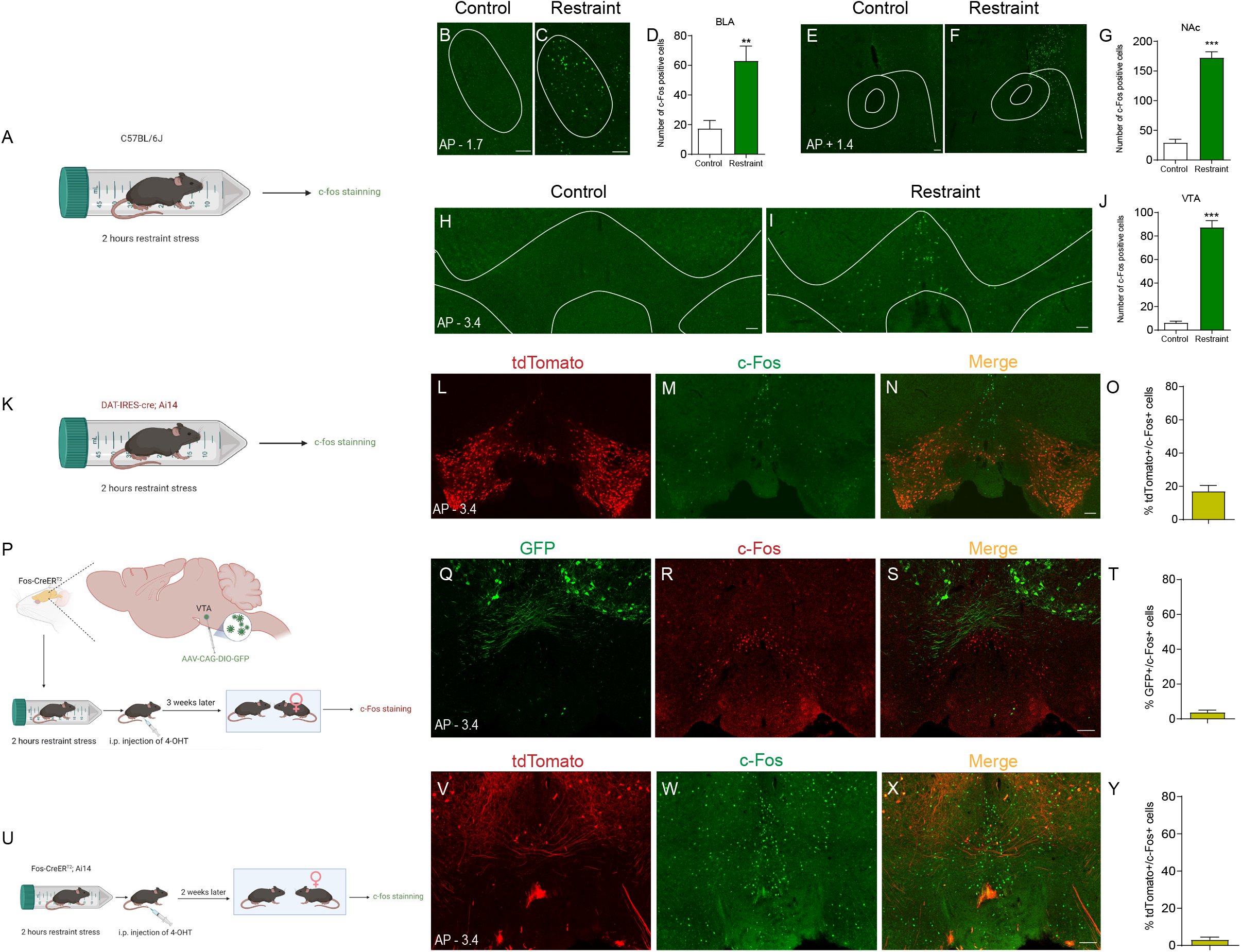
Positive and negative experiences activate distinct VTA neuron subpopulations. (A) Schematic showing the experimental procedure for C57BL/6J mice subject to restraint stress and subsequent c-Fos staining. Representative images showing c-Fos expression in BLA (B and C), NAc (E and F) and VTA (H and I). Statistical analysis showing restraint stress increased c-Fos expression in BLA (D, two-tailed unpaired t test, t_(10)_ =3.989, P = 0.0026), NAc (G, two-tailed unpaired t test, t_(10)_ = 12.23, P < 0.001) and VTA (J, two-tailed unpaired t test with Welch’s correction, t_(5.610)_ = 13.69, P < 0.001). (K) Schematic showing the experimental procedure for DAT-IRES-Cre;Ai14 mice subject to restraint stress and subsequent c-Fos staining. Representative images showing the tdTomato (L) and c-Fos (M) expression in VTA. (N) Images of L and M are merged. (O) Statistical analysis showing the percentage of c-Fos-positive VTA neurons that were labeled by tdTomato (17.04±3.513%, n=4). (P) Diagram illustrating virus injection in target areas and subsequent experiments. Representative images showing the GFP (Q) and c-Fos (R) expression in VTA. (S) Images of Q and R are merged. (T) Statistical analysis showing the percentage of c-Fos-positive VTA neurons that were labeled by GFP (3.765±1.285%, n=5). (U) Schematic showing the experimental procedure for Fos-CreER^T2^; Ai14 mice. Representative images showing the tdTomato (V) and c-Fos (W) expression in VTA. (X) Images of V and W are merged. (Y) Statistical analysis showing the percentage of c-Fos-positive VTA neurons that were labeled by tdTomato (3.098±1.412%, n=5). ***P<0.001. Data are presented as means +/− SEM. Annotation (AP): distance from the bregma (mm). Scale bars correspond to 100μm.

### GABAergic neurons activated by restraint stress inhibit the dopaminergic neurons responding to positive experience

The majority non-dopaminergic VTA cells are GABAergic neurons, which make up about 30% of the total neurons (Dobi et al, 2010). VTA GABAergic neurons have been recognized as potent mediators of reward and aversion, regulating behavioral outputs through projecting to distal brain regions or inhibiting local VTA dopaminergic neurons (van Zessen et al, 2012; Zhou et al, 2019; Bocklisch et al, 2013; Simmons et al, 2017). It has been previously reported that electric footshock activates VTA GABAergic neurons which inhibit local dopaminergic neurons (Dautan et al, 2016). In our study, the VTA neurons activated by restraint stress were mostly non-dopaminergic (Fig 6O), which suggested a possibility that the VTA non-dopaminergic cells responding to restraint stress were GABAergic neurons which inhibited the VTA dopaminergic neurons activated by positive experience, subsequently resulting in dysfunctions in reward-seeking behaviors.

To test this possibility, we first identified whether the VTA neurons responding to restraint stress were GABAergic using Vgat-ires-Cre mice. We injected AAV-DIO-mCherry into the VTA of Vgat-ires-Cre mice. As the Vesicular GABA Amino Acid Transporter (Vgat) is expressed by GABAergic neurons (Vong et al, 2011), the expression of mCherry would be induced in VTA GABAergic neurons. Then, we subjected these mice to 2 hours of restraint stress and performed c-Fos staining (Fig 7A). The result showed that most c-Fos-positive neurons were mCherry-labelled (79.49±3.652%, n=5) (Fig 7L-7H), suggesting that the activated VTA neurons are mostly GABAergic. We next performed in vivo extracellular recording to monitor the activity of VTA GABAergic neurons after restraint stress (Fig 7I) based on the electrophysiological criteria described in previous reports (Tan et al, 2012; Steffensen et al, 1998; Ko et al, 2018). Consistent with the histological results, restraint stress increased the number of spontaneous firing GABAergic neurons (Fig 7J), but did not influence their firing rate (Fig 7K), suggesting that the non-dopaminergic cells activated by restraint stress are probably GABAergic neurons. Then, we expressed hM3D selectively in VTA neurons responding to restraint stress by injecting AAV-DIO-hM3D-mCherry (or AAV-DIO-mCherry as control) into VTA of Fos-CreER^T2^ mice. We subjected these mice to 2 hours of restraint stress and immediately intraperitoneally injected 4-OHT to induce the expression of hM3D in activated VTA neurons. We performed in vivo extracellular recording to monitor the activity of VTA dopaminergic neurons in those mice exposed to conspecific females 45 min after CNO injection (Fig 7L). Reactivation of VTA neurons activated by restraint stress decreased the number (Fig 7M) and percentage of burst firing (Fig 7O) of VTA dopaminergic neurons responding to positive experience, but did not affect their firing rate (Fig 7N). Collectively, our data showed that restraint stress increased the activity of VTA GABAergic neurons which inhibited the activity of VTA dopaminergic neurons responding to positive experience, hence resulting in dysfunction of the reward processing circuit and subsequent depressive-like behaviors.

**Figure 7.**
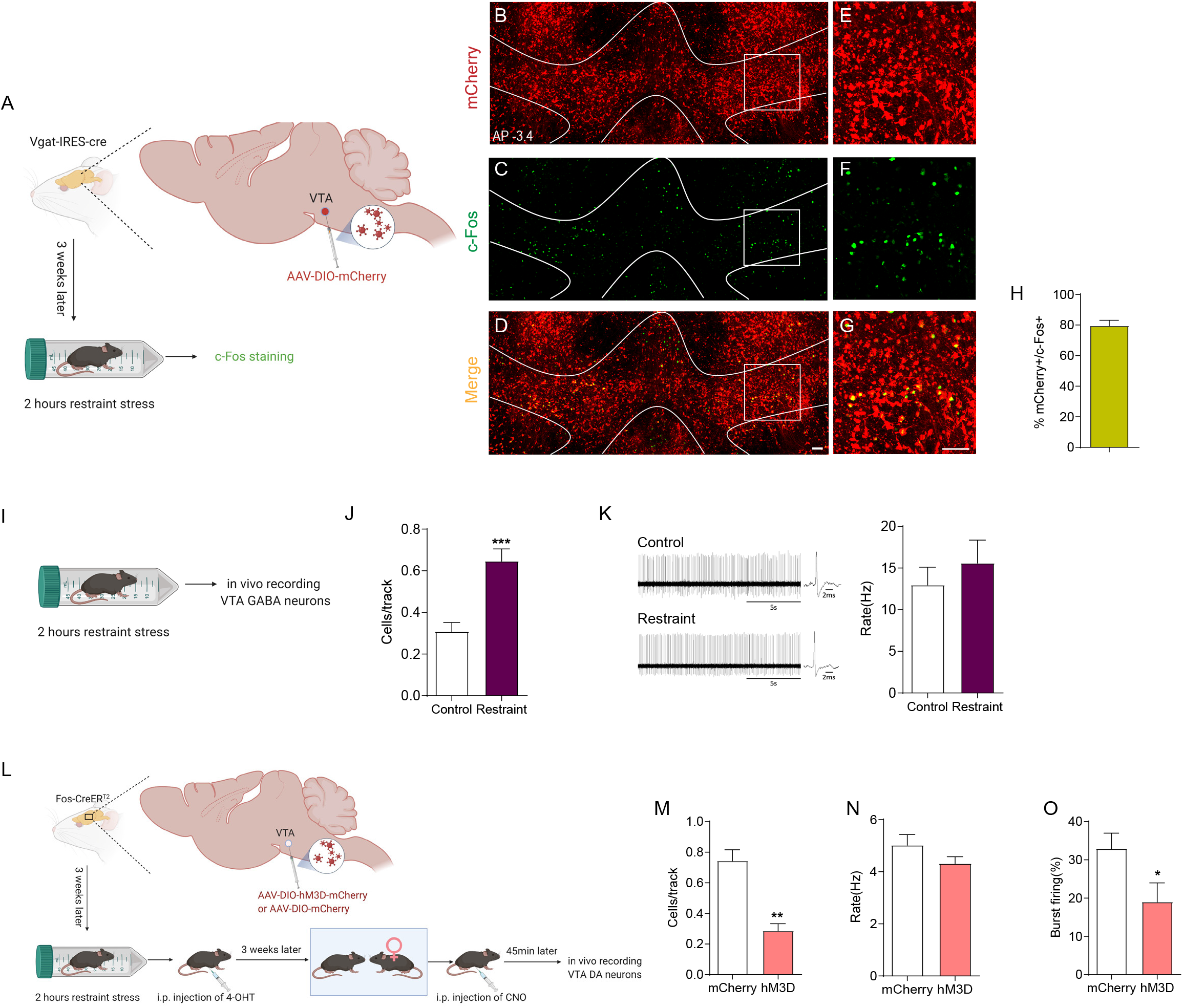
The VTA GABAergic neurons activated by restraint stress inhibit local VTA DA neurons responding to positive experience. (A) Diagram illustrating virus injection in target areas and subsequent experiments. Representative images showing the expression of mCherry (B, E) and c-Fos staining (C, F) in VTA. (D, G) Images are merged. (H) Statistical analysis showing the percentage of mCherry -labeled VTA neurons positive also for c-Fos (79.49±3.652%, n=5). (I) Schematic showing the experimental procedure for C57BL/6J mice subject to restraint stress and subsequent in vivo extracellular recording. (J) Restraint stress increased the number of spontaneously active GABA neurons per track in the VTA (two-tailed unpaired t test, t_(13)_ = 4.461, P=0.0006; Control n=7 mice, Restraint n=8 mice). (K) Left, representative extracellular voltage traces from VTA GABAergic neurons; Right, statistical result showing the average firing rate of the active GABAergic neurons (Mann-Whitney U test, P=0.8192; Control n=7 mice (13 neurons), Restraint n=8 mice (31 neurons)). (L) Diagram illustrating virus injection in target areas and subsequent experiments. Reactivation of VTA neurons previously activated by restraint stress inhibited the responsiveness of VTA dopaminergic neurons to sexual reward, decreasing the number of spontaneously active dopaminergic neurons per track in the VTA (M, two-tailed unpaired t test, t_(11)_ = 5.299, P=0.0003; mCherry n=6 mice, hM3D n=7 mice) and the percentage of burst firing (O, Mann-Whitney U test, P=0.0350; mCherry n=6 (40 neurons), hM3D n=7 mice (22 neurons)), but no change in the firing rate(N, Mann-Whitney U test, P=0.2281; mCherry n=6 (40 neurons), hM3D n=7 mice (22 neurons)). *P<0.05, ***P<0.001. Data are presented as means +/− SEM. Annotation (AP): distance from the bregma (mm). Scale bars correspond to 100μm.

## Discussion

In this study, we used anterograde and retrograde monosynaptic tracing, combining c-Fos immunofluorescence, to elucidate the architecture of the VTA-BLA-NAc circuit activated by sexual reward experience (positive experience). Projection-specific chemogenetic blockade of this circuit induced depression-like behaviors under normal conditions, and reactivation of VTA neurons activated by positive experience could ameliorate depression-like behaviors caused by chronic restraint stress. Furthermore, the subpopulation of VTA neurons responding to positive experience was mostly dopaminergic, while the VTA neurons activated by restraint stress belonged to anatomically distinct cell subpopulation in which non-dopaminergic neurons predominated and exerted inhibitory effect on the dopaminergic neurons activated by positive experience.

The mesocorticolimbic dopaminergic system originating from the VTA dopaminergic neurons which chiefly project to NAc, BLA, mPFC, and hippocampus, plays an essential role in reward, motivation, cognition, and aversion (Morales & Margolis, 2017; Fields et al, 2007). Its dysfunction has been implicated in many neuropsychiatric disorders including depression (Vrieze et al, 2013), bipolar disorder (Burdick et al, 2014), schizophrenia (Davis et al., 1991), and addiction (Koob & Le Moal, 2001; Loureiro M & Lüscher C, 2018). The VTA-NAc pathway has been extensively studied, which is critical for motivation and reward processing (de Jong et al, 2019; Saddoris et al, 2015; Mohebi et al, 2019), as well as addiction (Martínez-Rivera et al, 2017; Lüscher, 2016). The VTA dopaminergic neurons can facilitate or suppress target NAc neural activity not only directly through dopaminergic receptors residing on NAc neurons (Soares-Cunha et al, 2016; Pascoli et al, 2015), but also by regulating excitatory glutamatergic inputs to NAc neurons originating from BLA via presynaptic mechanisms (Stuber et al, 2011; Charara & Grace, 2003), integrating different inputs and turning them into action via outputs to ventral pallidum (Creed et al, 2016) and lateral hypothalamus (Luo et al, 2018; Maldonado-Irizarry et al, 1995). In our study, positive experience significantly increased active VTA, BLA and NAc neurons (Fig 1), and reactivation of VTA neurons activated by positive experience could enhance the neural activity of BLA and NAc (Fig 2A-2O), suggesting that BLA and NAc were target nuclei of VTA during positive experience. The results obtained from c-Fos staining experiment combined with retrograde tracing showed that almost all of BLA neurons activated by positive experience synapsed onto NAc (Fig 2V’’), while only few activated VTA neurons projected to NAc (Fig 2X’’), suggesting that reactivation of VTA neurons activated by positive experience increased the NAc neural activity probably through activating BLA neurons projecting to NAc. There is some evidence obtained from neuropharmacological experiments indicating that the VTA-BLA-NAc neural circuit is involved in modulating rewarding effects and motivated behaviors (Di Ciano & Everitt, 2004; Lintas et al, 2011), but direct anatomical evidence still lacks. We determined the putative neuronal circuit using the WGA-Cre transsynaptic tracing technology (Libbrecht et al, 2017; Hadpech et al, 2018), and the results showed that the mCherry labelled BLA neurons received inputs from VTA and sent projections to NAc (Fig 2P-2S), providing neuroanatomical evidence for the VTA-BLA-NAc circuit.

The prominent clinical feature of depression is anhedonia, and dysfunction of brain reward processing system has been implicated in neuropsychiatric disorders such as bipolar disorder (Caseras et al, 2013), schizophrenia (Strauss & Gold, 2012) and depression (Nestler & Carlezon, 2006; Pizzagalli et al, 2009). To determine whether dysfunction of the VTA-BLA-NAc circuit activated by positive experience induces depressive-like bahaviors, we labelled VTA or BLA neurons activated by positive experience with hM4D in Fos-CreER^T2^ mice and selectively inhibited the hM4D-labelled axon terminals projecting from VTA to BLA or from BLA to NAc through infusion of CNO into BLA or NAc, then performed FUST and SPT to evaluate the depression level. Our results showed that blocking the VTA-BLA or BLA-NAc pathways responding to positive experience induced depressive-like behaviors (Fig 3). The present medication treatments for depression, with the view that depression is a general brain dysfunction, take weeks to alleviate depressive symptomatology, while approximately 30% of patients get little improvement and are defined as having treatment-resistant depression (TRD) (Akil et al, 2018; Fogelson & Leuchter, 2017). More focused, targeted treatments that modulate specific brain networks or areas, such as deep brain stimulation (DBS), may prove to be promising approaches to help treatment-resistant patients (Kiening K & Sartorius A, 2013; Mayberg et al, 2005; Dandekar et al, 2018). A recent study by Steve Ramirez and his colleagues demonstrated that activating DG cells associated with positive memory could alleviate chronic stress-induced behavioral impairments (Ramirez et al, 2015). We therefore investigated whether directly activating the neural circuitry responsible for reward processing during positive experience could ameliorate depression-like behaviors induced by chronic restraint stress. We found that reactivating the VTA neurons activated by positive experience through CNO/hM3D system could ameliorate the depressive-like behaviors induced by chronic restraint stress (Fig 4).

The VTA dopaminergic neurons mediate a diverse array of functions associated with distinct axonal targets, including reward-related learning, goal-directed behavior, working memory, and decision making (Morales & Margolis, 2017; Björklund & Dunnett, 2007; Baimel et al, 2017). They exhibit rapid burst firing in response to rewarding stimuli or rewarding cues, subsequently sharply increasing dopamine release on their target brain regions (Watabe-Uchida et al, 2017; Pignatelli & Bonci, 2015; Lohani et al, 2018; Lavin et al, 2005). In our study, VTA neurons activated by positive experience were mostly dopaminergic, and their burst firing increased (Fig 5). Some studies have shown that the VTA dopaminergic neurons are also excited by aversive stimuli (Bromberg-Martin et al, 2010; Brischoux et al, 2009; Budygin et al, 2012), while there are paradoxical reports that they are inhibited by aversive events (Ungless et al, 2004; Matsumoto & Hikosaka, 2009). Recently, increasing evidence suggests that GABAergic neurons of the VTA, the majority of VTA non-dopaminergic neurons, are involved in mediating reward and aversion (Tan et al, 2012; van Zessen et al, 2012; Eshel et al, 2015). In our study, c-Fos staining revealed that restraint stress also induced neuron activation in VTA, BLA and NAc (Fig 6A-6J), but the activated VTA neurons were mostly non-dopaminergic (Fig 6K-6O). Consistent with a previous report that aversive stimulus activated VTA GABAergic neurons (Tan et al, 2012), our histological and electrophysiological results showed that restraint stress increased the number of spontaneously active GABAergic neurons in VTA (Fig 7A-7K), indicating the activated non-dopaminergic neurons were probably GABAergic. Our data suggested that the VTA may process reward and aversive stimuli through dopaminergic and GABAergic neurons respectively. Given the difference in VTA neuron types responding to restraint stress and sexual reward, how does restraint stress influence the performance of reward-seeking behaviors (Fig 4). Our results revealed the distinct anatomical locations of these two VTA neuron subpopulations (Fig 6P-6Y), but we noticed that the restraint stress-responding neurons sent fibers to the anatomic location of positive experience-responding neurons, indicating that there may be synaptic connections between these distinct neuron subpopulations. It is generally accepted that the VTA GABAergic neurons promote aversive behaviors through inhibiting VTA dopaminergic neurons (Tan et al, 2012; Bocklisch et al, 2013), prompting us to determine whether VTA neurons activated by restraint stress inhibited the responsiveness of VTA dopaminergic neurons to positive experience. Positive experience increased the number of active dopaminergic neurons and the percentage of burst firing neurons (Fig 5N-5Q), which would be inhibited by reactivation of VTA neurons activated by restraint stress (Fig 7L-7O). Our data was similar and consistent with a previous study by Ruud van Zessen and his colleagues, which showed that nonspecific of activation of the VTA GABA neurons reduced the excitability of neighboring VTA dopaminergic neurons in vitro (van Zessen et al, 2012), while our work specifically targeted the VTA neuron subpopulation responding to restraint stress and provided electrophysiological evidence in vivo. Repeatedly activating the VTA neuron subpopulation responding to restraint stress may reduce the excitability and responsiveness of the VTA neuron subpopulation processing reward stimuli, which may be the underlying mechanism by which chronic stress induces impaired reward-seeking behaviors. This needs to be clarified in future studies.

In conclusion, we show that the VTA-BLA-NAc circuit, whose dysfunction would induce depressive-like behaviors, is activated by positive experience, and reactivation of the VTA neurons activated by positive experience can ameliorate the depressive-like behaviors induced by chronic restraint stress. Furthermore, we reveal that reactivation of VTA neuron subpopulation responding to restraint stress inhibits the activity of the VTA neurons responding to positive experience, and that this may be the mechanism by which chronic restraint stress may induce anhedonia, the core feature of depression.

## Materials and Methods

### Animals

C57BL/6 mice were purchased from the Pengyue Laboratory of China, and transgenic mice (DAT-IRES-Cre, 006660; Ai14, 007908; Fos-CreER^T2^, 021882; Vgat-IRES-Cre, 016962) were purchased from the Jackson Laboratories. The protocols of the animal studies were approved by the Institutional Animal Care and Use Committee of Binzhou Medical University Hospital and performed in compliance with the National Institutes of Health Guide for the Care and Use of Laboratory Animals. Efforts were made to minimize animal suffering and the number of animals used.

### AAV virus

AAV2/9-hSyn-DIO-hm3D(Gq)-mCherry (in the main text referred to as AAV-DIO-hm3D-mCherry), AAV2/9-hSyn-DIO-hm4D(Gi)-mCherry (in the main text referred to as AAV-DIO-hm4D-mCherry), AAV2/9-hsyn-DIO-mCherry (in the main text referred to as AAV-DIO-mCherry), AAV2/9-hsyn-WGA-Cre-T2A-ZsGreen (in the main text referred to as AAV-hSyn-WGA-Cre-T2A-ZsGreen), AAV2-CAG-Retro-GFP (in the main text referred to as AAV-Retro-GFP) were purchased from Shanghai Hanheng Biotechnology, China.

### Drugs

Clozapine-N-oxide (CNO) was purchased from Sigma, dissolved in saline at a concentration of 0.5 mg/ml and diluted in saline to a final concentration of 0.03 mg/ml (hM3D) for intraperitoneal injection. For intra-BLA and NAc microinjection (hM4D), CNO was diluted in saline to a final concentration of 3mM. 4-Hydroxytamoxifen (4-OHT) was purchased from Sigma and dissolved in ethanol. Corn oil (Sigma) was added to the 4-OHT solution, which was shaken and mixed at 37°C, put in a fume hood for ethanol volatilization to a final concentration of 5 mg/ml and stored at −20°C with protection from light.

### Stereotaxic surgery

For microinjection of AAV virus, mice were anaesthetized and mounted onto a stereotaxic frame (KOPF, USA). AAV virus was injected with a glass pipette using an infusion pump (Micro 4, WPI, USA). Viruses were injected bilaterally into target brain areas using the following coordinates: VTA, AP=-3.30 mm, ML= ± 0.50 mm, DV=-4.20 mm, from bregma; flow rate of 0.10 μl/min and total volume of 1.00 μl (0.50 μl/side); NAc, AP=+1.30 mm, ML= ± 0.50 mm, DV=-4.80 mm, from bregma; flow rate of 0.10 μl/min and total volume of 0.60 μl (0.30 μl/side); BLA, AP=-1.80 mm, ML= ± 3.00 mm, DV=-4.80 mm from bregma, flow rate of 0.10 μl/min and total volume of 0.60 μl (0.30 μl/side). An additional 5 min was allowed for diffusion and prevention of backflow. Behavioral tests or in vivo extracellular recordings were conducted 21 days after AAV injection.

For BLA and NAc cannula implantation, adult male C57BL/6 mice were anaesthetized and mounted onto a stereotaxic frame (KOPF, US). The skull surface was coated with Kerr phosphoric acid gel etchant (Kerr USA). First, a bilateral guide cannula was inserted into the BLA (coordinates: 1.7mm posterior to bregma, 3.0 mm lateral to midline, and 3.8 mm ventral to dorsal) or NAc (coordinates: 1.4 mm anterior to bregma, 0.5 mm lateral to midline, and 3.8 mm ventral to dorsal). Then, adhesive (GLUMA, Germany) was applied onto the skull and cannula surface, and Resina fluida (Filtek Z350 XT 3M, USA) was brushed on top with light curing for 45 seconds using a VALO curing light (Ultradent Products). Finally, dental cement was used to seal the cannula, and a dummy cannula was inserted into the guide cannula to maintain unobstructed cannula. After surgery, animals were individually housed and then allowed to recover for 21 days with daily handling. Mice were conscious, unrestrained and freely moving in their home cages during the microinjections. On the experimental day, a 33-G stainless-steel injector connected to a 5-μl syringe was inserted into the guide cannula and extended 1 mm beyond the tip of the guide cannula. CNO (900pmol/0.3μl/side) or vehicle was infused bilaterally over 2.5 min. The injector tips were held in place for an additional 5 min after the end of infusion to avoid backflow through the needle track. Behavioral tests were performed 45 min after microinjections.

### Electrophysiology

Mice were anaesthetized with 4% chloral hydrate (400 mg/kg, intraperitoneally). The core body temperature was sustained at 37°C via a thermostatically controlled heating pad during the whole process. Nine electrode tracks (100μm interval, grid pattern) were placed in the VTA (coordinates: 3.3 to 3.5 mm posterior to the bregma, 0.3 to 0.5 mm lateral to the midline, and 3.5 to 5.0 mm below the brain surface). Putative VTA dopamine and GABA neurons were identified using established electrophysiological criteria (dopamine neurons: unfiltered waveform duration >2.2 ms overall, start-to-trough waveform duration ≥1.1 ms with high-pass filter, firing rate range from 0.5 to 10 Hz (Tan et al, 2012; Grace & Bunney, 1983; Ungless & Grace, 2012); GABA neurons: waveform duration <1 ms, firing rate range from 5 to 60 Hz (Tan et al, 2012; Steffensen et al, 1998; Ko et al, 2018)). The recording time for each neuron was over 3 min. Three parameters of VTA dopamine neuron activity were measured: (1). the number of spontaneously active dopamine neurons per track; (2). average firing rate; and (3) average percentage of burst firing, which is defined as the occurrence of two consecutive spikes with an inter-spike interval <80 ms, and the termination of a burst defined as two spikes with an inter-spike interval >160 ms (Grace & Bunney, 1983; Ungless & Grace, 2012).

### Immunohistochemistry and cell counting

Mice were anaesthetized and transcardially perfused with cold PBS and 4% paraformaldehyde (PFA) sequentially. Mouse brains were maintained in 4% PFA at 4°C overnight and then dehydrated in 30% sucrose for 2 days. Coronal sections (40μm) containing the target brain region were obtained using a freezing microtome (Leica, CM1950). The sections were incubated in blocking buffer (10% normal goat serum, 0.3% Triton X-100) for 1 hour at room temperature and then incubated with one or two primary antibodies overnight at 4°C, followed by rinsing in PBS buffer and secondary antibody incubation for 4 hours at room temperature. The sections were mounted with Gold antifade reagent containing DAPI (Invitrogen, Thermo Fisher Scientific). An Olympus FV10 microscope was used to capture images. Primary antibodies: c-Fos (CST, #2250) and tyrosine hydroxylase (TH) (ImmunoStar, #22941). The numbers of c-Fos, TH, GFP, mCherry and tdTomato positive cells were counted bilaterally by experimenters who were blind to the treatments. For each brain, the counting criteria for interested brain regions is described as follow: BLA, 3-4 slice, (every ninth section, from 0.94mm to 2.18 posterior to bregma); NAc, 3-4 slice, (every fourth section, from 1.78mm to 1.10 anterior to bregma); mPFC, 3-4 slice, (every third section, from 1.98mm to 1.54 mm anterior to bregma); VTA, IPN, and SN, 3-4 slice, (every third section, from 3.16mm to 3.64 posterior to bregma); DG, 3-4 slice, (every sixth section, from 1.46mm to 2.54 posterior to bregma). For each brain, the number of immunopositive cells of interested brain regions of per section was calculated by dividing the total number of immunopositive cells in all selected sections by the number of selected sections. For co-immuno experiments, using the percentage of TH positive of tdTomato labelled cells in VTA (Fig 5H) as an example, we divided the total number of co-immuno cells (TH+/tdTomato+) in VTA of all selected slices by the total number of tdTomato positive cells in VTA of all selected slices.

### Behavioral procedures

#### Positive experience (sex reward) and neutral experience

On the experimental day, both groups mice were moved to a behavior room with dim lighting conditions and housed individually. After 4 hours of habituation, positive experience group male mice were exposed to oestrous female mice for 2 hours. For neutral experience, male mice were housed with fake toy mice.

#### Acute and chronic restraint stress

Mice were moved to a behavior room on the experimental day and housed individually. After 4 hours of habituation, mice were exposed to two hours of restraint stress. Control mice were housed individually in the behavior room for 6 hours without any treatment. For chronic restraint stress treatment, stress group mice were subjected to 2 hours of restraint for 15 consecutive days.

#### Female urine sniffing test (FUST)

The female urine sniffing test was performed as previously described (Malkesman et al, 2010). On the experimental day, mice were transferred to a dimly lit behavior room at least 4 hours before beginning the experiment. The test procedure was as follows: 1. 3-min exposure to the cotton tip dipped in water; 2. 45-min interval; 3. 3-min exposure to the cotton tip infused with fresh urine from female mice in the oestrus phase. The duration of female urine sniffing time was scored.

#### Sucrose preference test (SPT)

Mice were habituated to drinking water from two bottles for one week before beginning testing. On the experimental day, water was deprived for three hours, and then two bottles were introduced (1% sucrose and water). Mice had free choice of either drinking 1% sucrose solution or water for 2 hours after lights off during the dark cycle. Sucrose and water consumption were determined by measuring the weight changes. Sucrose preference was calculated as the ratio of the mass of sucrose consumed versus the total mass of sucrose and water consumed during the test.

#### Forced swim test (FST)

The Plexiglas cylinder used for this test was 25 cm high and 10 cm in diameter. Each mouse was placed in a Plexiglas cylinder with water at a 15 cm depth (24°C) for 6 min, which was recorded by a camera directly above. The latency to immobility at the first 2 min and the duration of immobility during the last 4 min were measured. Immobility was defined as no movements except those that maintain their head above water for respiration.

#### Locomotor

This test was performed in SuperFlex open field cages (40 × 40 × 30 cm, Omnitech Electronics Inc., Columbus, OH), and mice were allowed 30 min free exploration under illuminated conditions. The total distance travelled was quantified using Fusion software (Omnitech Electronics Inc., Columbus, OH).

### Statistical analyses

Statistical analysis was performed with graphpad prism software. Results are presented as mean ±standard error of mean (S.E.M.). Shapiro–Wilk test was used to test the normality and equal variance assumptions. For normally distributed data, two-tailed t tests were used to assess differences between two experimental groups with equal variance. For a two-sample comparison of means with unequal variances, two-tailed t tests with Welch’s correction were used. For non-normally distributed data, Mann–Whitney U tests were performed to compare two groups. For multiple groups, two-way ANOVAs followed by Tukey’s multiple comparisons test were used. P< 0.05 was considered statistically significant.

### Data availability

This study includes no data deposited in external repositories.

## Acknowledgment

This work was supported by the following grants: National Natural Science Foundation of China (81901380 and 81500930), the Natural Science Foundation of Shandong Province (ZR2017BC047 and ZR2014HQ014).

## Author Contributions

H.Y. and B.L. conceived this study and designed the experiments. H.Y., B.L., and M.C. wrote the manuscript. L.S., J.Y., M.C., J.W., F.S., W.W., D.W., D.L., Z.X., and C.Q. performed the experiments and analyzed the data.

